# Dynamic regulation of HYL1 provides new insights into its multifaceted role in Arabidopsis

**DOI:** 10.1101/396861

**Authors:** Prakash Kumar Bhagat, Deepanjali Verma, Raghuram Badmi, Alok Krishna Sinha

## Abstract

MicroRNAs (miRNAs) are 21 to 24 nucleotide non-coding RNAs that regulate gene expression. Biogenesis of miRNAs is fine-tuned by specialized microprocessor complex, the regulation of which is being continuously understood. Recruitment of HYL1 to the microprocessor complex is crucial for accurate primary-miRNA (pri-miRNA) processing and accumulation of mature miRNA in *Arabidopsis thaliana*. HYL1 is a double-stranded RNA binding protein also termed as DRB1, has two double-stranded RNA binding domain at N-terminal and a highly disordered C-terminal region. Also, the biological activity of HYL1 is dynamically regulated through transition from hyperphosphorylation to hypophosphorylation state. HYL1 is known to be phosphorylated by a MAP kinase MPK3 and SnRK2. However, the precise role of its phosphorylation are still unknown. Recently, the stability of HYL1 protein has been shown to be regulated by an unknown protease X. However, the identity of the protease and its molecular mechanisms are poorly understood. Here, we describe, three functionally important facets of HYL1, which provide a better picture of its association with molecular processes. First, we identified a conserved MPK3 phosphorylation site on HYL1 and its possible role in the miRNA biogenesis. Secondly, the C-terminal region of HYL1 displays tendencies to bind dsDNA. Lastly, the role of C-terminal region of HYL1 in the regulation of its protein stability and the regulation of miRNA biogenesis is documented. We show the unexplored role of C-terminal and hypothesize the novel functions of HYL1 in addition to miRNA biogenesis. We anticipate that the data presented in this study, will open a new dimension of understanding the role of double stranded RNA binding proteins in diverse biological processes of plants and animal.

## Introduction

MicroRNAs (miRNAs) are endogenous small regulatory RNA, known to regulate diverse functions during growth and development in both plants and animals (Reinhart et al., 2000; Voinnet et al., 2009; Rogers and Chen, 2013). In Arabidopsis, the biogenesis of mature miRNAs is tightly regulated by microprocessor complex consisting of HYPONASTIC LEAVES 1/Double-stranded RNA Binding protein (HYL1/DRB1) as one of the essential factor (Kurihara et al., 2006; Manavella et al., 2012). HYL1 along with Dicer-like 1 (DCL1) and SERRATE (SE) binds to the primary-and/or precursor-miRNA (pri-/pre-miRNA) structures and executes the accurate miRNA processing during miRNA biogenesis (Yang et al., 2006; Lobbes et al., 2006; Song et al., 2007; Dong et al., 2008). MicroRNA biogenesis is very important in regulating various developmental and environmental responses of the plant, as *hyl1* mutant displays pleiotropic phenotype (Lu and Fedoroff, 2000). Post-translational regulation of HYL1 has recently gained much attention and it is known that the functions of HYL1 is regulated by the phosphorylation/dephosphorylation cycle (Manavella et al., 2012; Raghuram et.al. 2015; Yan et al., 2017; Su et al., 2017). It is known that HYL1/DRB1 are phosphorylated by mitogen-activated protein kinase-3 (MPK3) in Arabidopsis and rice (Raghuram et.al. 2015). MPK3 phosphorylation site on HYL1 as well as its biological significance is still not clear. It is also known that AtCPL1/2 (C-Terminal Phosphatase Like 1/2) dephosphorylates HYL1 and regulates its functions in miRNA biogenesis (Manavella et al., 2012). Constitutive photomorphogenic 1 (COP1) protects HYL1 light-dark dependent manner (Cho et al., 2014). However, HYL1 phosphorylation and its role on HYL1 stability, its implications on light-dark regulation are still unknown. HYL1, as other dsRNA binding proteins possesses two double–stranded RNA binding domains (dsRBD) at its amino-(N-) terminal, which is responsible for interaction with miRNA transcript. However, the carboxyl-(C-) terminal region of HYL1 is being understood as of less importance in HYL1’s functions (Wu et al., 2007; Yang et al., 2010; Liu et al., 2013; Baranauskė et al., 2015).

Here, we investigate the multifaceted role of HYL1/DRB1 by (i) domain fragmentation analysis, (ii) deciphering phosphorylation site by exploiting naturally occurring mutations in HYL1 orthologs and (iii) conditional subcellular localisation of HYL1. Our work also provides caution to the researchers who use the HYL1 antibody (Agrisera#AS06136) and its possible limitations to study the dynamically regulated HYL1 like proteins.

## Results and Discussion

### MPK3 phosphorylates HYL1 at conserved sites

To identify MPK3 phosphorylation site/s on HYL1, we generated the N-terminal (1-170 AA) and C-terminal (171-419 AA) constructs of HYL1, thus separating the RNA-binding domains fragment and the disordered C-terminal fragment (Fig. 1a). *In-vitro* phosphorylation assay using bacterially expressed AtHYL1N (1-170 AA), AtHYL1C (171-419 AA) and AtHYL1FL (Full-Length) indicated that AtMPK3 phosphorylates both N-terminal and C-terminal fragments as well as the full-length AtHYL1 protein (Fig. 1b). To further investigate the interaction specificity, we generated the fragmented constructs of the two RNA-binding domains (RBDs) at the N-terminal of HYL1 (AtHYL1R1 and AtHYL1R2). Yeast two-hybrid AD constructs harbouring AtHYL1FL, AtHYL1R1, AtHYL1R2, AtHYL1N and AtHYL1C were used to determine the interaction specificity with AtMPK3. As shown in Fig. 1c, AtMPK3 interacts with all the analysed AtHYL1 fragments including the full-length, thus confirming our observations from *in-vitro* phosphorylation assay (Fig. 1b). Extending these observations to the previously known HYL1 interactors, CPL1/2 (Manavella et al., 2012) and SE (Baranauskė et al., 2015), and to analyse the observations in the context of the microprocessor complex, we performed a yeast two-hybrid assay using CPL1/2 and SE as baits. In addition to their observed interactions with full length HYL1 (Manavella et al., 2012; Fig. 1c), we found that CPL1/2 interacts with both N-and C-terminal regions of HYL1 whereas SE specifically interacts with N-terminal region (Fig 1d and 1e). These observations clearly indicate that AtMPK3 phosphorylation sites as well as CPL1/2 dephosphorylation sites are present on both N-and C-terminal regions of AtHYL1.

**Figure 1.**
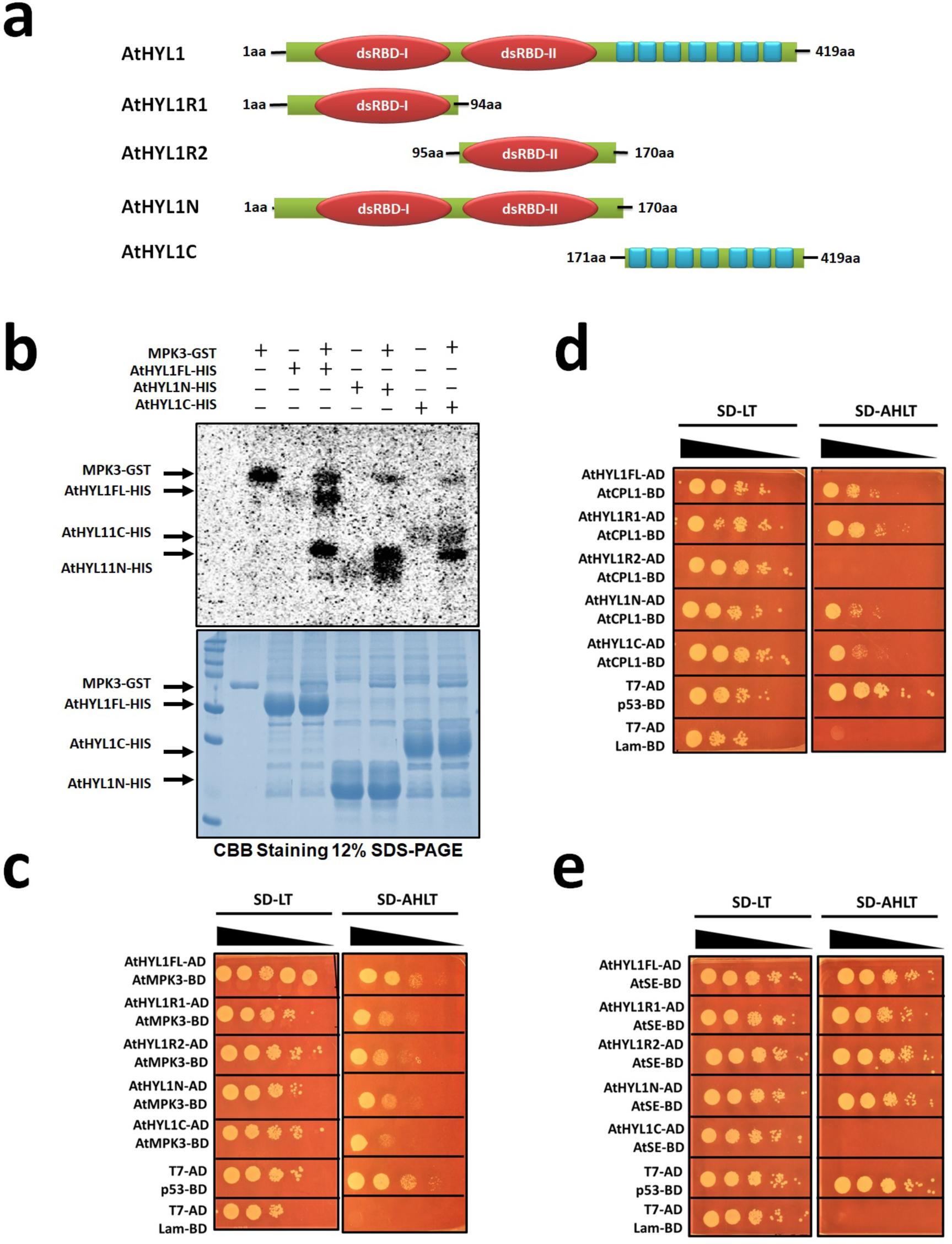
HYL1 is phosphorylated at multiple sites by MPK3. a, Diagrammatic representation of AtHYL1 protein and its domain organisation. The N-terminal and C-terminal half used for protein expression, *in-vitro* experiments and Y2H are indicated by amino acid numbers. **b,** *in-vitro* phosphorylation assay showing the phosphorylation of HYL1 full length (AtHYL1FL), N-terminal (AtHYL1N) and C-terminal (AtHYL1C) regions by AtMPK3. **c to e,** Yeast two-hybrid assay showing the interaction of different versions of AtHYL1 (AtHYL1FL, AtHYL1N, AtHYL1C, AtHYL1R1 – first dsRBD and AtHYL1R2 – second dsRBD) **c,** with AtMPK3 **d,** with CPL1 and **e,** with SE. Images were taken after growing the yeast on respective medium at 28°C for 3 to 4 days.

Sequence analysis and comparison of the closest AtHYL1 orthologs in different Arabidopsis and other species revealed two interesting insights: (i) the two RBDs in the N-terminal region are highly conserved in all the analysed species whereas the extended C-terminal is not (Fig 2 and Supp Fig. 1a) and (ii) the first RBD in all the analysed sequences contain a conserved ‘TP’ motif (Fig. 2, Supp Fig. 1a), that is a putative MAPK phosphorylation site (Sharrocks et al., 2000). This observation is in agreement with the conserved interactions between HYL1/DRB1 and MPK3 in Arabidopsis and rice (Raghuram et al., 2015) and interaction of CPL1/2 with HYL1. This indicated the high likelihood of the RBD1 ‘TP’ motif to be a conserved MPK3 phosphorylation site present in distantly related species (Supp Fig. 1b).

**Figure 2.**
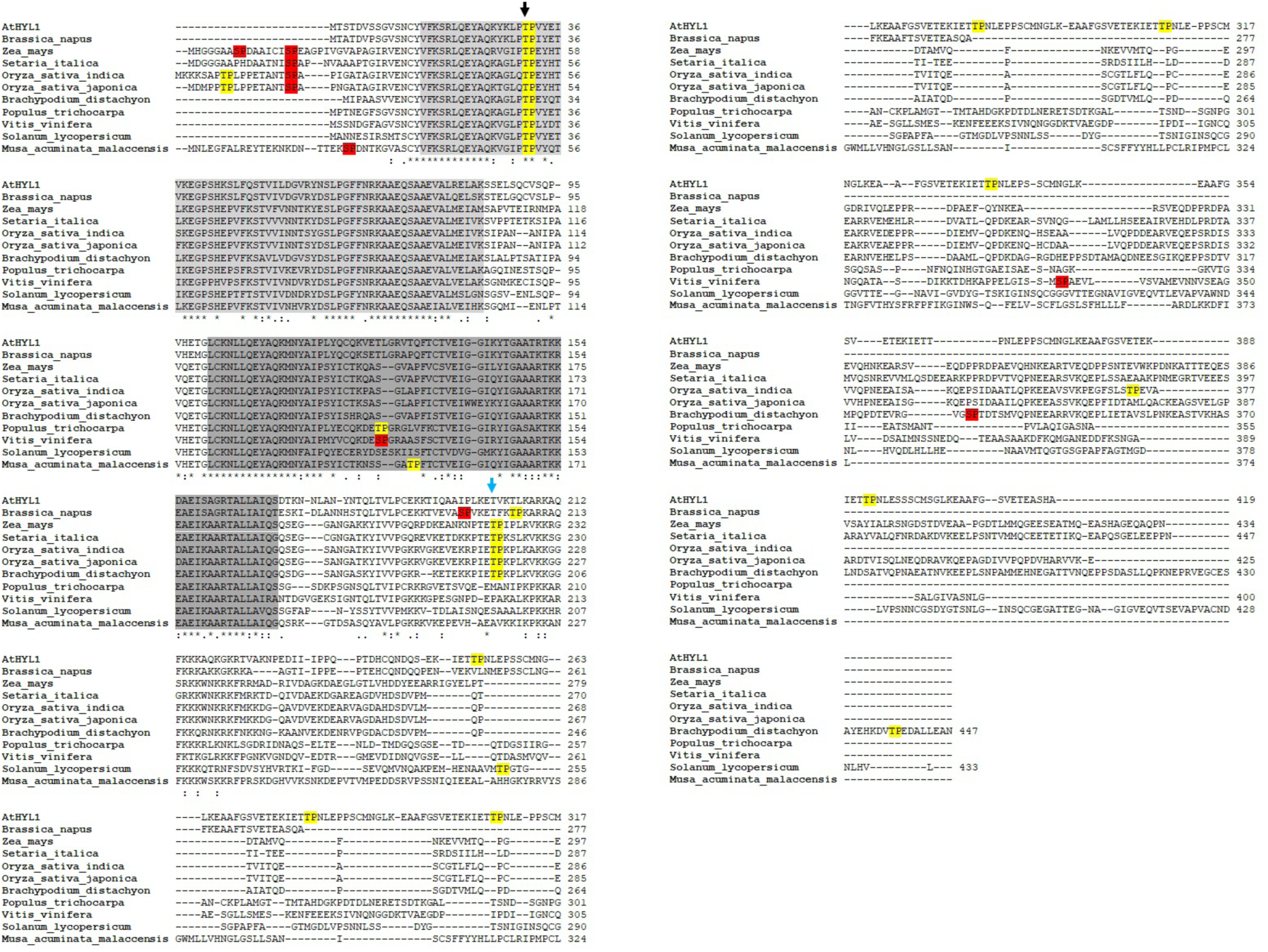
Putative MAP kinase site is evolutionarily conserved in plant kingdom. The Amino acid sequence alignment of full-length HYL1/DRB1 proteins from *Arabidopsis thaliana* and other plant species as described in the method section. The domains are highlighted by light grey (dsRBD-I) and dark grey (dsRBD-II). The putative MAP kinase SP and TP motifs are highlighted by red and yellow respectively. The evolutionarily conserved TP motif present at dsRBD-I in N-terminal is indicated by black arrow whereas, newly evolved TP motif present in the *A. alpine* and all other monocots at C-terminal is indicated by sky blue arrow.

To investigate these aspects in more detail, we extended the observations to the monocot crop plant rice. We observed that one of the OsDRBs, OsDRB1-4 lacks the conserved ‘TP’ motif in its RBD1 (Supp Fig. 2a) and that T (Threonine) is replaced by L (Leucine). We exploited this naturally occurring mutation to investigate the phosphorylation site of MPK3 in RBD1. We generated the N-and C-terminal fragmented constructs of OsDRB1-1, OsDRB1-2 and OsDRB1-4 for bacterial expression and used the fragmented proteins in an *in-vitro* phosphorylation assay by OsMPK3. Excitingly, we observed the phosphorylation in both N-and C-terminal fragments of OsDRB1-1 and OsDRB1-2 and only in the C-terminal fragment of OsDRB1-4, but not in the N-terminal fragment (Fig. 3). These observations combined with yeast two-hybrid results (Fig. 1b and Fig. 1c) point out that the ‘TP’ motif in the RBD1 (T31 of AtHYL1) is the MPK3 phosphorylation site conserved in distantly related species.

**Figure 3.**
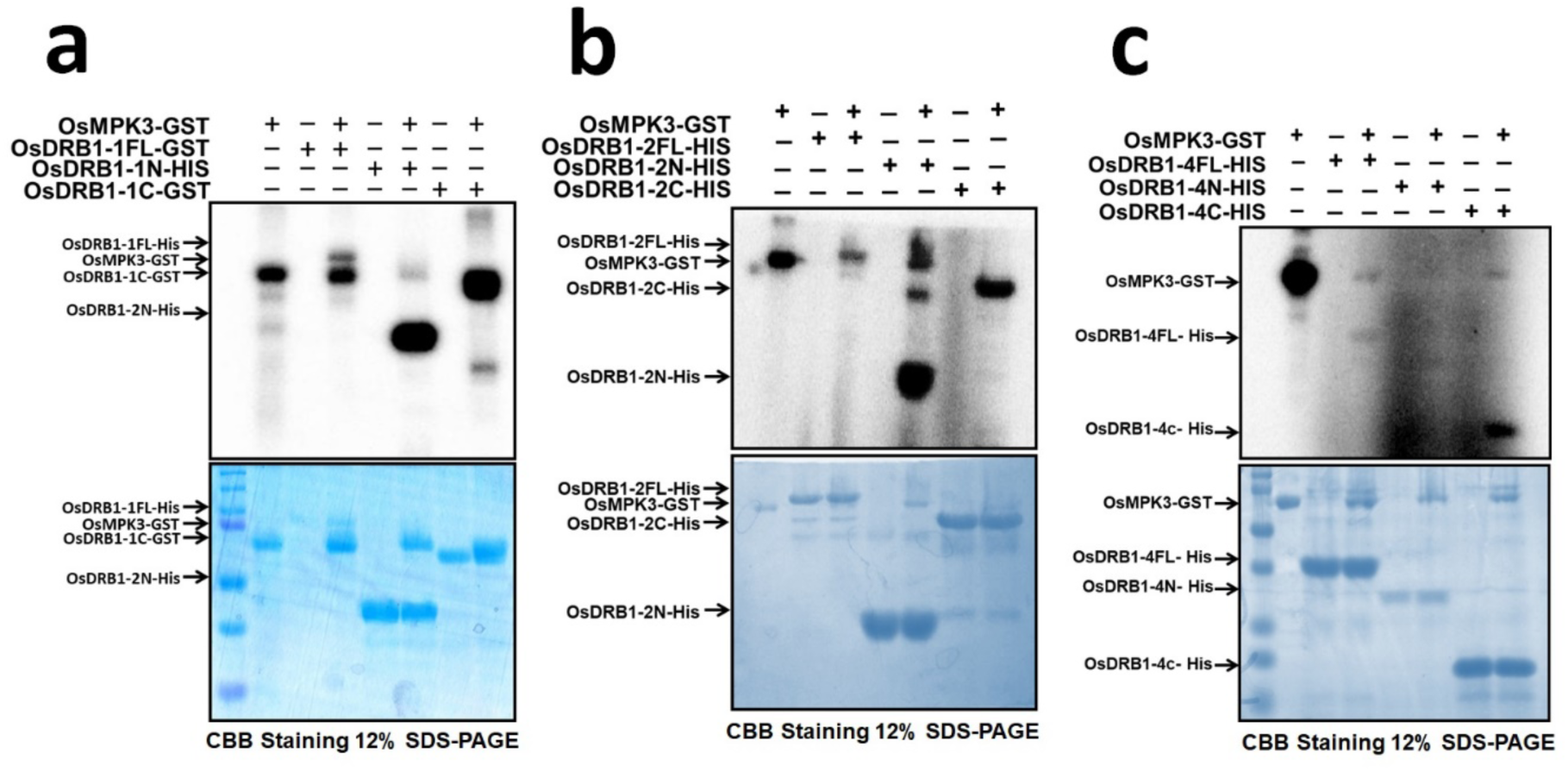
The conserved threonine at dsRBD-I is phosphorylated by MPK3. a, *In-vitro* phosphorylation assay showing the phosphorylation of full length and truncated proteins of OsDRB1 **b,** OsDRB1-2 and **c,** OsDRB1-4 by OsMPK3. The upper images are the autoradiograph and lower are CBB staining of respective gels. The positions of phosphorylated protein are indicated by arrows.

### The DNA binding properties of HYL1

#### The disordered and non-conserved C-terminal region of HYL1

Earlier studies pointed out that the C-terminal repeat region has minimal biological functions (Wu et al., 2007). Our observation that MPK3 interacts and phosphorylates the non-conserved C-terminal region (Fig. 1b, Fig. 1c and Fig. 3) led us to question this assumption. Sequence analysis revealed the presence of non-conserved putative MAPK phosphorylation ‘TP’ motif in few of the analysed sequences but not all (Fig 2 and Supp Fig. 1a). In the sequences with the C-terminal ‘TP’ motifs, these motifs were present in different regions of the C-terminal domain (Fig 2 and Supp Fig. 1a). We hypothesize that the six C-terminal ‘TP’ motifs present in AtHYL1 is responsible for its hyper-and hypo-phosphorylation status. Thus phosphorylation at multiple sites by MPK3 or other kinases, may accumulate the hyperphosphorylated isoform of HYL1 as reported earlier (Manavella et al., 2012). Dephosphorylation at any of these threonine residues by CPL1 may result into biologically active, hypophosphorylated HYL1. However, the absence of C-terminal repeats and ‘TP’ motifs in other plant species and lesser number of 28 aa repeats in other members of Arabidopsis might hint to the evolutionary answers. It can be hypothesised that, during the evolution of these proteins, the C-terminal region has gone a very strict selection pressure by a series of elimination of repeats, one by one, as we can see their variable numbers in other Arabidopsis species (SI Fig. 1). Other possible explanation can be that the repeats region is duplicated several times in *A. thaliana*.

We asked, if there are any other proteins which are regulated by the phosphorylation-dephosphorylation cycle at repetitive regions by MAP kinases and phosphatases? We found a classical example of the CTD (C-terminal domain) repeats in RNA polymerase II (RNA pol II) in yeast, animals and plants. The heptad repeats (Y^1^S^2^P^3^T^4^S^5^P^6^S^7^) is present at the carboxyl terminal of the RNA pol II. The dynamic phosphorylation of two serine residues S^2^ and S^5^ has been observed in CTD repeats by MAP kinases ERK1/2 in human, yeast and *Xenopus laevis* Oocyte. In plants, MPK3 phosphorylates RNA pol II and regulates its activity (Bellier et al., 1997; Bonnet et al., 1999; Prelich et al., 2009; Glover-cutter et al., 2009; Zhang et al., 2016). The dephosphorylation of CTD by CPL1 is associated with transcription initiation by interaction of RNA pol II at the promoter with TATA binding proteins. Phosphorylation of CTD by MPK3 leads to disruption of preinitiation complex and transition to formation of elongation complex with other proteins. Hence the reversible phosphorylation regulated by MAP kinase and CPL1 regulates the transcription process. These reports combined with the HYL1 literature (Manavella et al., 2012; Raghuram et al., 2014) points out that MPK3 and CPL1 are the regulators of HYL1 and RNA pol II, the two major regulators of RNA metabolism. The sharing of common regulators by these two major proteins, participating in the RNA metabolism may have some homologous role that is dependent on reversible phosphorylation. To analyse this proposed function, the carboxyl terminal of AtRPB1 (RNA polymerase II larger subunit 1) and AtHYL1 was further analysed for natural disorder regions in these proteins. The composition of amino acids in a protein makes naturally ordered/disordered regions on the protein surface which are proposed to participate in the protein –protein and protein – nucleic acid interactions (Jamsheer et. al., 2018). The *in-silico* analysis revealed that both proteins, HYL1 and RNA pol II have ordered regions in their N– terminals, however, C-terminal is highly disorder in both HYL1 (AtHYL1C) and AtRPB1 (1254 to 1839 amino acids includes CTD repeats, RBP1C) (SI Fig.3). We mutated the phosphorylation sites *in-silico* and analysed the C-terminal region of RNA pol II for its ordered/disordered properties. The mutated phosphorylation sites(phosphor-mimetic) display the changes in orderedness than the native versions. The dynamic phosphorylation sites are concentrated at the disordered regions at C-terminal which is true in case of AtRPB1 and AtHYL1. As the C-terminal repeats of both RNA binding proteins are phosphorylated by MPK3, we presume that AtHYL1C might function similar to the CTD repeats of AtRBP1.

#### C-terminal region enhances DNA binding properties of HYL1

The *in-silico* observations led us to hypothesize the comparable functions of the HYL1 C-terminal domain to that of RNA pol II CTD. HYL1 is known to recognise the dsRNA hairpin like structure that describes its specificity for secondary structure of RNA rather than the nucleotide specificity (Lu and Fedoroff, 2000). We argued that if the recognition is only dependent on the structure and not the sequence, then AtHYL1C might recognise the double-stranded DNA (dsDNA) hairpin loop. Such properties have been previously described and studied to function in a variety of processes (Hudson et al., 2014; Cassiday et al., 2002). To test this hypothesis, chemically synthesized hairpin forming single stranded DNA oligos (Fig. 4a) were used for the electrophoretic mobility shift assay (EMSA) using AtHYL1-FL (full-length) and AtHYL1N protein. Surprisingly, we observed that AtHYL1-FL binds strongly to dsDNA hairpin loop than that of AtHYL1N (Fig. 4b). This observation is in contrast to the previous reports, where the authors have shown the positive EMSA interaction of HYL1 with dsRNA but not with dsDNA (Lu and Fedroff., 2000). Interestingly, most of the studies since the first report used dsRNA and there is no evidence for HYL1 interaction with dsDNA.

**Figure 4.**
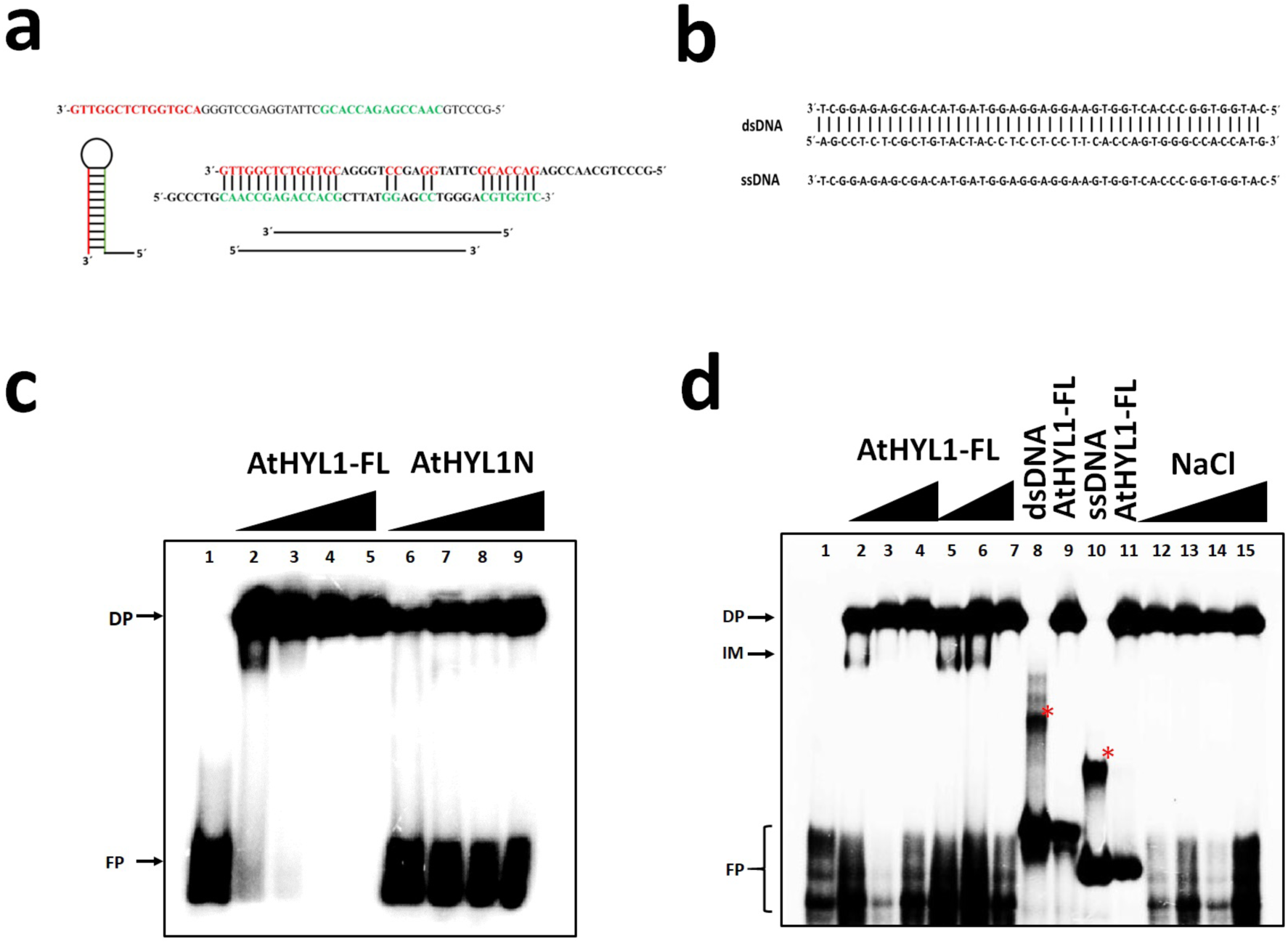
The carboxyl terminal has the propensity to regulate the HYL1 interaction with nucleic acid *in-vitro*. a, The diagrammatic representation of chemically synthesised ssDNA and its dsDNA hairpin loop and possible dimer. **b,** dsDNA and ssDNA. **c,** Gel mobility shift assay showing the interaction of full length AtHYL1 and truncated AtHYL1N proteins with dsDNA hairpin loop in 5 % polyacrylamide gel prepared in TBE. Lane 1, the labelled probe alone, lane 2-5, the probe was incubated with increasing concentration of HYL1FL protein. **d,** showing the gel mobility shift of AtHYL1 full length proteins at higher gel percentage (10% PAGE). Lane 1, is the free probe alone, lane 2-4 and 5-7, the increasing concentration of HYL1FL carried out from two separate protein batch with dsDNA hairpin loop, lane 8, double-stranded DNA (dsDNA) probe alone, lane9, dsDNA with HYL1FL, lane 10, ssDNA probe, lane 11, ssDNA with HYL1FL protein, lane 12-15, dsDNA hair pin loop with HYL1FL protein with increasing concentrations of salt (NaCl). The free probe (FP) are at the bottom and DNA-protein (DP) complex and intermediate (IM) are indicated by arrow. Red asterisk indicates the intermediate forms of dsDNA and ssDNA without proteins.

The DNA binding ability of the full-length HYL1 protein suggests a possible role of its C-terminal repetitive regions (Fig. 4b). Judging by the amount of unbound free probe, we observe that the deletion of C-terminal domain in HYL1 protein drastically reduced its binding ability with dsDNA hair pin loop as compared the HYL1-FL (Fig. 4b). This clearly suggests that C-terminal disorder regions have the propensity for recognition for dsRNA/dsDNA together with the N-terminal double stranded RNA binding domains (dsRBD). This observation can be further substantiated by the previous reports where N-terminal of HYL1 was shown to be sufficient for complimenting the *hyl1* mutant phenotype and resumes the miRNA biogenesis (Wu et al.,2007). Therefore, even in the absence of C-terminal, the AtHYL1N binds the hairpin loop (Yang et al., 2010) but the strength of binding is low as compare to full length AtHYL1FL, even at higher concentrations of protein (Fig. 4c).

While analysing the HYL1 interaction with dsDNA hairpin loop by EMSA, we observed a DNA-protein complex lower to the intense band at lower concentration of protein (Fig. 4b, lane 2). When we analysed the same experiment on higher percent gel, this complex was further resolved very clearly (Fig 4d, lane 2-7). We initially, thought it to be an intermediate complex. Dimerization analysis of the ssDNA also showed the probability for double – stranded DNA (dsDNA) formation. Thus to analyse the interaction with other forms of DNA i.e dsDNA and ssDNA we again performed the EMSA with full length HYL1 protein (Fig. 4b). Result clearly showed the interaction of HYL1 with both forms of DNA (Fig. 4d, lane 8 −11). These interactions were further strengthened by increasing the concentration of salt (NaCl) which established that these interactions are very strong (Fig. 4d, lane 12 -15). The interaction of HYL1 with dsDNA and other forms lead us to suggest that HYL1 may have unknown multiple functions in nucleic acid metabolism and its regulation. One possible role could be its participation during the transcription of miRNA or mRNA primary transcript by interaction with DNA and other proteins at promoter regions along with the transcription initiation complex and RNA pol II as discussed above.

### HYL1 stability and localisation

#### The unknown protease X is a Trypsin-like protease

Recent studies show that an unknown protease regulates HYL1 stability by proteolytic degradation in the cytosol and that COP1 protects HYL1 from degradation (Cho et al., 2014) However, the identity of this protease and its implication on miRNA biogenesis is still unknown. Trypsin digestion has long been used to map the functional domains of proteins (Lorence et al., 1988 and Zvaritch et al., 1990). Trypsin is an endopeptidase and has higher specificity for peptide cleavage on C-terminal side of arginine (R) and lysine (K). The presence of acidic amino acid at either side of target residues slow down the rate of hydrolysis and merely presence of proline (P) on carboxyl side abolishes the cleavage. Our *in-silico* analysis of HYL1 protein sequence identified potential trypsin cleavage sites at the junction of the AtHYL1N and AtHYL1C region (bipartite NLS) and in every 28aa repeat region of AtHYL1C (Supp. Fig. 4). Further, the molecular weight of HYL1 cleaved product is known to be around 25 kDa (Cho et al., 2014), which matches exactly with *in-silico* prediction. We therefore, performed time-course trypsin digestion of bacterially expressed AtHYL1FL-His (Fig. 5a). Very interestingly, we observed two bands with time-dependent increase in intensity, which migrate at about 25 kDa. Time-course digestion reactions with AtHYL1N-His were unaltered, pointing out the stability of AtHYL1N from trypsin digestion. The lower band of AtHYL1FL-His digestion products matches the size of AtHYL1N-His, suggesting the upper band is AtHYL1C-His. Further, we tested the degradation of AtHYL1FL-His using protein extract from wildtype *A. thalinana* Col-0 tissue and AtHYL1N-His as reaction controls. As seen in Fig. 5b, we observed time-course dependent digestion of AtHYL1FL-His as observed by the increase in intensity of the band at 25 kDa, with matching size of unaltered AtHYL1N-His (Fig. 5b). These observations suggest that the unknown protease X (Cho et al., 2014) that cleaves AtHYL1 can presumably be a trypsin-like protease (TLP). To further validate our observations, western blotting was performed using anti-HYL1 antibody (Agrisera) using the trypsin digestion reaction products, again by using AtHYL1N-His as reaction controls. To our surprise, we observed time-dependent decrease in intensities of both AtHYL1FL-His and its degraded products at 25 kDa (Fig. 5c). The absence of signal from AtHYL1N-His sample wells (Fig. 5c) prompted us to uncover that the anti-HYL1 antibody was specific to the C-terminal region of HYL1 (https://www.agrisera.com/en/artiklar/hyl1-hyponastic-leave-phenotype-ds-rna-binding-protein.html) and does not recognise AtHYL1N-His. Furthermore, time-course trypsin digestion of upto five hours confirmed that longer incubation leads to decrease in intensities of the 25 kDa bands pointing out the digestion of the C-terminal region, where the potential trypsin cleavage sites were identified in each 28aa repeats (Fig. 5d). These results support, HYL1 is degraded by TLP into N-and C-terminal cleaved products and that C-terminal region is further degraded.

**Figure 5:**
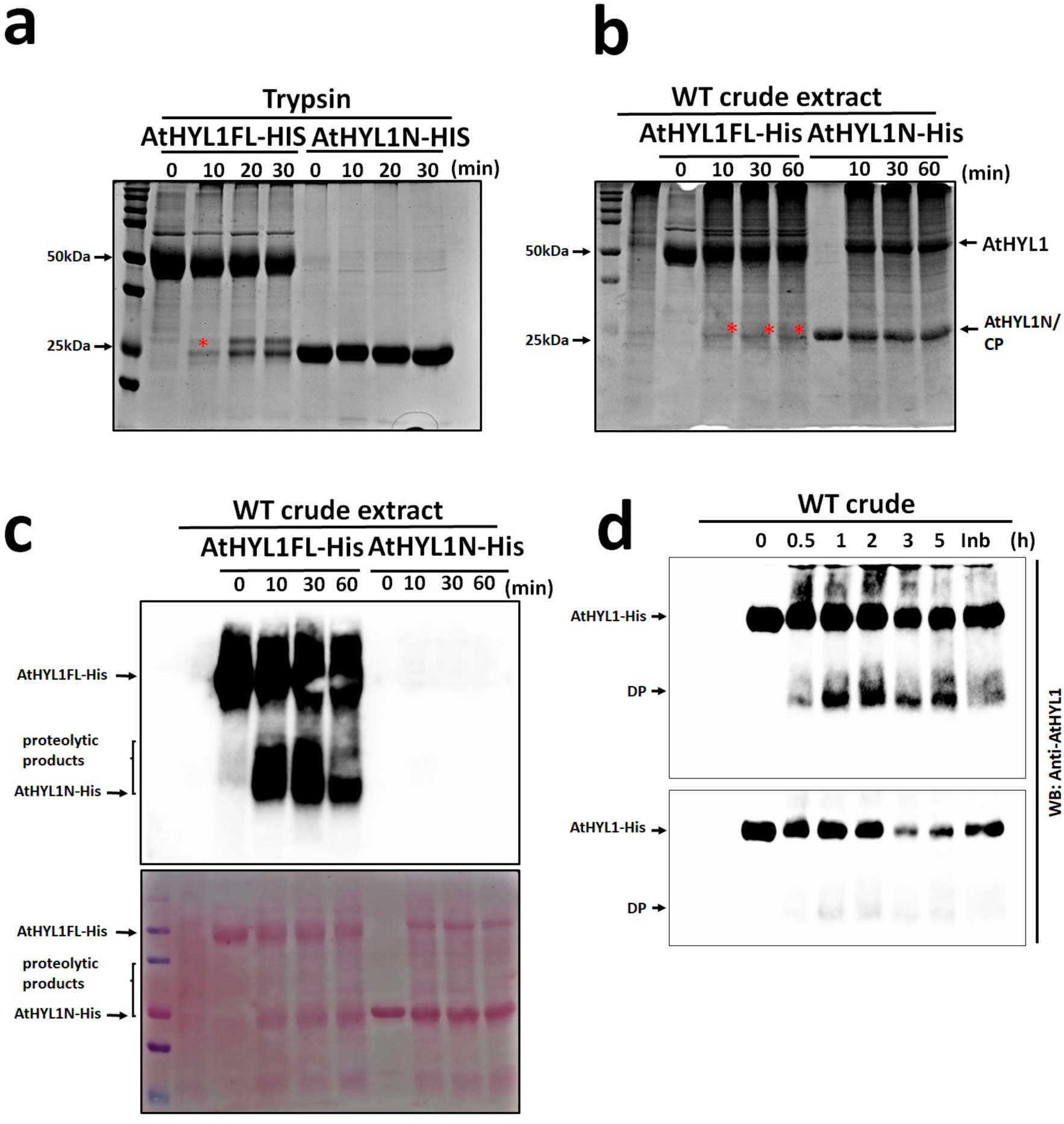
AtHYL1 is cleaved by protease at multiple sites in the carboxyl terminal repeats. a, Protease sensitivity assay showing the cleavage of AtHYL1 full length and truncated AtHYL1N by trypsin. The reactions were incubated for indicated times periods followed by SDS-PAGE and gel was stained by CBB. **b**, in-vitro protein degradation assay of bacterially purified AtHYL1 full length and truncated AtHYL1N incubated with crude protein extract from wild type *A. thaliana* (*Col-0*) for indicated time periods followed by CBB staining of SDS-PAGE. **c**, experiment was performed as in b, followed by western blotting with anti-AtHYL1 antibody. Lower image is ponceau staining of the membrane. **d**, in-vivo degradation of AtHYL1FL with crude extract (Col-0) with extended periods of incubation (0.5 hrs to 5 hrs) followed by western blotting using anti-AtHYL1. The upper image represents the longer exposure and lower shorter exposure during the western blotting detection. DP represents degraded products and PPI indicates the plant protease inhibitor cocktail.

As the TLP is known to be present in the cytoplasm (Cho et al., 2014), we hypothesized that cleavage by TLP will influence the observed AtHYL1 localisation, depending upon the terminal of GFP-tag. As the most probable TLP cleavage sites are present in NLS (Supp Fig. 4: R-222, K-228 and K-250), cleavage at one of these residues should result into disruption of its nuclear localization, rendering the fluorescence outside of nucleus. We generated two versions of GFP-tagged HYL1 constructs – GFP-AtHYL1 and AtHYL1-GFP and performed tobacco agro infiltration to observe HYL1 localisations. Excitingly, the two constructs yielded two different localisation profiles of AtHYL1 (Fig. 6a and 6b). The N-terminal GFP-tagged GFP-AtHYL1 was localised in the nucleus and extra nuclear compartments like cytoplasm and membrane (Fig. 6a). Whereas, the C-terminal GFP-tagged AtHYL1-GFP was exclusively localised into the nucleus (Fig. 6b). This difference in localisation patterns explain two things: (i) the cytoplasmic TLP protease cleaves AtHYL1 so that the bipartite NLS is retained with the C-terminal region of AtHYL1 (Fig. 6b), meaning that it is highly likely that the cleavage site of TLP is R-222, K-228 (Supp fig. 4), (ii) the N-terminal GFP-tagged GFP-AtHYL1 is stable in the extranuclear region (Fig. 6a) as it does not possess any TLP cleavage sites (Supp fig. 4).

**Figure 6:**
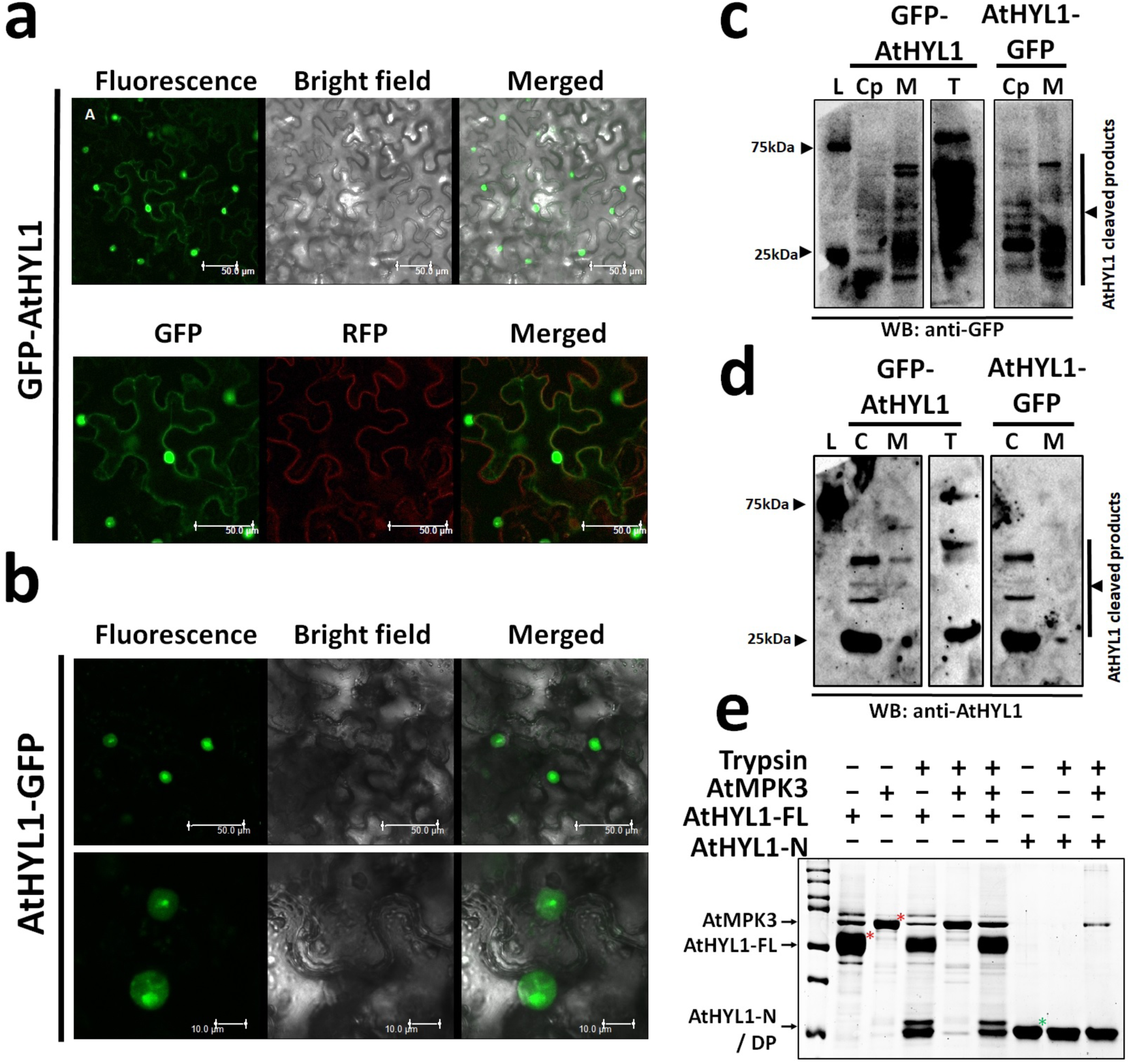
AtHYL1 localization and stability is regulated by its C-terminal regions. Subcellular localization of HYL1 was monitored by tagging the protein with GFP either at amino terminal (GFP-AtHYL1) (a) or at carboxyl terminal (AtHYL1-GFP) GFP-AtHYL1 (b) in the *N. benthamiana* leaves under confocal laser scanning microscopy. The scale bar is indicated in each images. Membrane marker tagged with RFP was used as positive control. The subcellular enriched fractions from above experiments were further used for western blotting with anti-GFP (c) and anti-AtHYL1 antibody. The C, M and T represents cytoplasmic, membrane and total crude extract respectively. (d). The protease sensitivity test was performed after in-vitro phosphorylation of AtHYL1 by AtMPK3 with cold ATP followed by trypsin digestion. Red asterisk indicates AtMPK3 (upper) and AtHYL1 (lower). Green asterisk represents truncated AtHYL1N.

To obtain further insights into AtHYL1 degradation by TLP, we expressed AtHYL1FL protein GFP tagged either at N-or C-terminal ends, and transiently expressed in *Nicotiana benthamiana* as above. Proteins were extracted from the different cellular compartments including enriched cytoplasmic and membrane fractions along with total protein. These enriched fractions were immunoblotted with anti-GFP and anti-HYL1.

Immunoblotting using fractionated proteins confirmed the localisation of AtHYL1 to cytoplasm and membrane providing further insights (Fig. 6c and 6d). Most of the extra nuclear AtHYL1 protein was present in the form of cleaved products. Results clearly showed the expression of HYL1 at about 70 kDa and 55 kDa present in the total crude extract from GFP-HYL1. Additionally, there were other bands from 26 kDa to 55 kDa and appeared as smear with longer exposure time. Immunoblotting analysis of cytoplasmic HYL1 protein using anti-GFP, did not show any signal when protein was tagged at amino-terminal (GFP-HYL1) (Fig. 6c). However, using the anti-AtHYL1 antibody showed an intense band at 25 kDa resembling truncated C-terminal of HYL1 (HYL1C) along with the other low intense bands which are likely the proteolytic cleaved products (Fig. 6d, lane 2). The membrane fraction showed an intense band of the truncated HYL1 (GFP-HYL1N) at about 55 kDa with anti-GFP along with the degraded products with lower molecular weight (Fig. 6c, lane 3). Blotting with anti-AtHYL1 did not show any intense band suggesting that only N-terminal is relocated to membrane (Fig. 6d, lane 3). When protein was tagged at the C-terminal with GFP showed an intense band at 25 kDa, presumably the truncated GFP after protease cleavage and three bands in the range of 25 kDa to 37 kDa. These are the different proteolytic bands from C-terminal regions (Fig. 6c, lane 6) which was further confirmed by using anti-AtHYL1 (Fig. 6d lane 6). Based on these results, we concluded that after the cleavage by TLP, primarily at NLS, the N-terminal is sorted to the membrane and the C-terminal is quickly degraded in the cytosol. The membrane localized truncated GFP-HYL1 (GFP-HYL1N) may be further transported into nucleus as reported earlier (Wu et al., 2007).

The importance of unique C-terminal with bipartite NLS and 28 amino acid repeats indicates its biological importance in the regulation of HYL1 localization and stability. The stable protein finally regulates the miRNA biogenesis and its rate of processing by sorting its functional domain. To test the role of MPK3 phosphorylation at C-terminal repeats, AtHYL1FL-His protein was incubated with AtMPK3 in an *in-vitro* kinase reaction before the proteolytic cleavage by trypsin. The result shows that MPK3 phosphorylation does not alter the proteolytic cleavage by trypsin (Fig. 6e).

#### Interaction with MPK3 overrides light-dark transition dependent localisation of HYL1

To investigate the influence of MPK3 phosphorylation on localisation patterns of AtHYL1 *in-vivo*, we co-expressed the GFP-AtHYL1 and AtMPK3-HA in the *N. benthamiana* leaves. To address the observation of light-dark transition dependent localisation (Achkar et al., 2018), we incubated the agro infiltrated plants either in light or in dark. Surprisingly, the localisation of MPK3 was restricted to the nucleus and no fluorescence was observed from cytoplasm and the membrane (Fig. 6a, 6b and Fig. 7). Specifically, the fluorescence was prominent from the nuclear bodies, presumably nuclear dicing or D bodies (Manavella et al., 2012). This suggests that MPK3 phosphorylation of HYL1 can override the light-dark transition dependent localisation as observed before (Achkar et al., 2018) These observations point out to a yet unknown mechanism of HYL1 regulation and portrays MPK3 as a major player in COP1 dependent HYL1 stability. It can be speculated that MPK3 overexpression retains COP1 in the cytoplasm, thus inhibiting TLP and protecting HYL1 from degradation. However, further experiments in this direction will reveal many interesting insights and uncover the under lying regulatory mechanism of diverse function of HYL1 protein.

**Figure 7.**
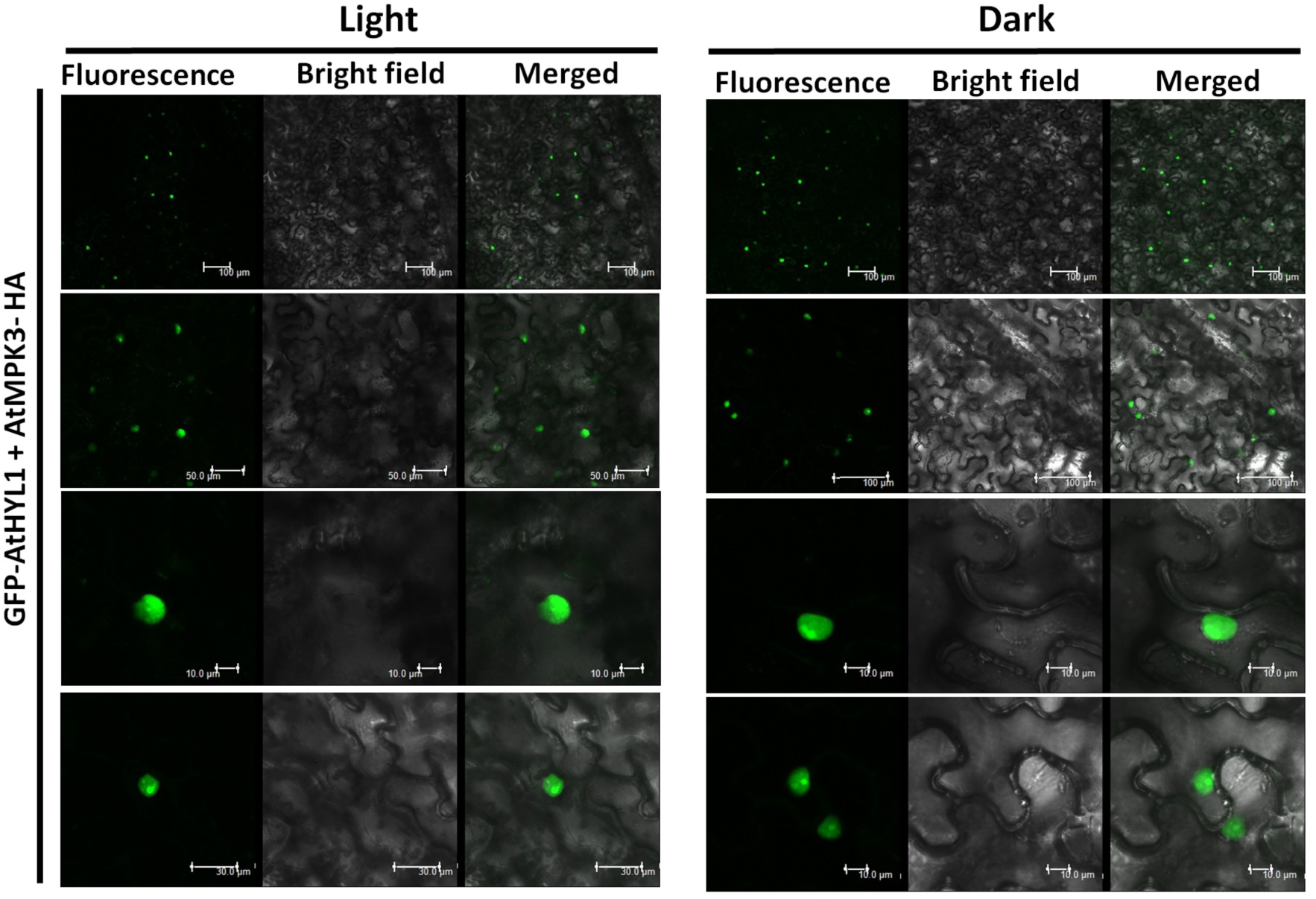
AtHYL1 phosphorylation by AtMPK3 regulates its localization and stability in-vivo. The subcellular localization of GFP-AtHYL1 was monitored in the presence of AtMPK3-HA protein in *N. benthamiana* leaves in response to light and dark. Agrobacterium harbouring above constructs were co-infiltrated and the localization of GFP-AtHYL1 was monitored in leave samples harvested from plants keep at light or dark for 8 to 10 hours, under confocal microscope. Scale bar is indicated in each images.

## CONCLUSIONS

1. MPK3 phosphorylation site in N-terminal region of HYL1 is conserved in distantly related species.
2. The C-terminal region of HYL1 is disordered, non-conserved in other species and enhances the DNA-binding ability of HYL1
3. HYL1 also possess dsDNA binding property.
4. Trypsin-like protease (TLP) is responsible for the cytoplasmic degradation of HYL1 and its regulation during light-dark transition
5. MPK3 overexpression renders nuclear localisation of HYL1

## OUTSTANDING QUESTIONS

1. What is the evolutionary significance of N-terminal HYL1 phosphorylation by MPK3?
2. Does HYL1 C-terminal region has minimal functions and insignificant in the context of evolution?
3. What is the identity of Trypsin-like protease (TLP) and the gene it encodes for it?
4. How does MPK3 overexpression influence the nuclear localisation of HYL1?

## MATERIALS AND METHODS

### Plant growth conditions

The *Arabidopsis thaliana Col-0* wild type seeds were surface sterilized and plated on half MS plate and incubated at 4°C for 2 to 3 days for stratification. The plates were further incubated at 22°C in the growth room with long day conditions. The rice (*Oryza sativa L. indica* cultivar group var. Pusa Basmati1) was grown at 28°C with 16 h light/ 8 h dark oscillation in a greenhouse at the National Institute of Plant Genome Research, New Delhi or in growth chamber (SCILAB instrument, Taiwan, China). The 7 to 10 days old seedling were harvested for RNA isolation and cDNA preparation.

### Yeast two-hybrid assay

The full length CDS of AtMPK3, AtSE, AtCPL1 and AtHYL1 along with the deletion fragments were amplified from the above prepared cDNA using the primer sets (SI Table1) by Phusion DNA-polymerase (NEB, USA) in – frame with pGADT7 (AD) and pGBKT7 (BD) (Clontech). Protein-protein interactions were performed according to manufacturer’s protocols. Briefly, the AD and BD constructs were co-transformed in yeast competent cells (Y2H gold) prepared and performed according to the G-bioscience (Fast – yeast transformation kit) and shredded on double dropout synthetic define(SD) medium lacking leucine and tryptophan (SD-LT) then interaction was monitored on Quadruple dropout medium lacking adenine, histidine, leucine and tryptophan (SD-AHLT) by incubating at 28-30°C. Whenever the interaction was measured with MPK3, 10-15 mM 3-amino-1,2,4-triazole (3-AT) was added in the medium. The picture was captured regularly from 2-4 days after serial dilution of respective yeast cells.

### *In-vitro* phosphorylation assay

To performed the *in-vitro* phosphorylation assay, the full length CDS of AtMPK3, OsMPK3 and OsDRB1-1 were cloned in – frame with pGEXT42 vector. The AtHYL1, OsDRB1-2 and OsDRB1-4 were cloned in the pET series vectors (pET21c/pET28a) along with the all deletions constructs using the primer sets (SI Table1). After DNA sequencing, all constructs were transformed in *Escherichia coli* BL21 strain. The proteins were purified using the Ni-NTA and GST-beads. The in-vitro phosphorylation assay was performed according to the previously described (Raghuram et.al., 2014). Briefly, the purified proteins and kinase were incubated in the kinase reaction buffer (25 mM Tris/Cl, pH 7.5, 5 mM MgCl2,25 mM ATP, 1 mM EGTA, 1 mM DTT, 5 µCi of γ-^32^P–ATP) at 30°C for 30 minutes. The reaction was terminated by addition of SDS (sodium dodecyl sulfate)-sample loading buffer followed by heat denaturation at 95°C for 5 minutes. Samples were separated on 10-15 % SDS– polyacrylamide gel electrophoresis (SDS-PAGE). The phosphorylation was detected and Coomassie Brilliant Blue (CBB) stained using phosphor imager, Typhoon (GE Healthcare, Life Sciences).

### Electrophoretic Mobility Shift Assay (EMSA)

The binding of AtHYL1-His and AtHYL1N-His was performed with chemically synthesized 50 nucleotides single – stranded DNA (ssDNA) (3´-GTTGGCTCTGGTGCAGGGTCCGAGGTATTCGCACCAGAGCCAACGTCCCG-5´). The ssDNA was labelled with γ^32^P-ATP at 30°C for 30 minutes by PNK-T4 kinase (NEB). The labelled ssDNA was purified and denatured at 95°C then allowed for formation of ds-DNA hairpin loop to mimic the dsRNA hairpin loop by subsequent cooling gradually at room temperature. The EMSA was performed as previously described (Kshirsagar et.al.,2017, Lorence et al, 1988 and Zvaritch et al, 1990). Briefly, increasing concentration of HYL1FL and HYL1N was added in the EMSA buffer containing10 mM Tris-HCl (pH 7.5), 350 mM KCl, 2.5 mM EDTA, 10% glycerol, 5 mM DTT, and 10 mM MgCl2 along with poly-dIdC at room temperature for 30 minutes. The samples were resolved on 5% to 8% of native polyacrylamide gels prepared and electrophoresed in 0.5% TBE (tris-boric acid and EDTA). The DNA-protein complex is visualized by phosphor imagers Typhoon. Similarly, EMSA was carried out with double – stranded DNA and ssDNA (SI table).

### *In-vitro* Protein sensitivity assay

The protease sensitivity assay using bacterially purified AtHYL1 and AtHYL1N terminal was performed according to previously described (Reddy et al., 2001; Yamamoto *et.al.*, 2014). Briefly, the 4-5µg proteins were partially digested in the digestion buffer (5 mM MgCl2, 100 mM KCl in 25 mM HEPES-KOH, pH 7.4) with trypsin (Sigma) at 37°C for indicated time. Reactions were stopped by addition of SDS-loading dye followed by heating at 95°C for 5 minutes. The samples were loaded on 12-15% SDS-PAGE and protein bands were visualized by Coomassie Brilliant Blue (CBB) staining. The western blotting of the same experiment set was further analysed by anti-AtHYL1 antibody (AS06136, Agrisera) according to Singh and Sinha. Briefly, after SDS-PAGE, the separated proteins were transferred to nitrocellulose blotting membrane (10600016, GE Healthcare life science). The membrane was blocked with 5% skimmed milk in TBS-T buffer (TBS0.1% and Tween 20) at room temperature for 2 hours, followed by incubation with primary antibody anti-AtHYL1 (1:1000) over night at cold room. The following day, membrane was washed with TBS-T buffer and incubated with goat anti-rabbit IgG secondary antibody HRP conjugate (31463, Thermo Fisher scientific, USA) with 1: 10,000 dilutions in blocking solution at room temperature for 1 to 2 hours. The membrane was washed same as above 3 to 4 times followed by washing with water. The membrane was developed using clarity^TM^ western ECL substrate (170-5061, Biorad).

### *In–vitro* protein degradation assay

The in-vitro protein degradation assay was performed as described previously (Cho *et al.*, 2014) with slightly modifications. Briefly, 2 to 5 microgram bacterially purified AtHYL1 full length and AtHYL1N with his tagged proteins were incubated with the total crude extract from Arabidopsis thaliana wild type for indicated time periods at 37°C. The samples were harvested and reaction was stopped by adding SDS-sample loading dye, heated at 95°C for 5 minutes and samples were loaded on 12-15% SDS-PAGE analysis. The protein gels were proceeded for both Coomassie Brilliant Blue (CBB) staining as well as immunoblotting using anti-AtHYL1 antibody as described above.

### Protein subcellular localization

The localization of full length HYL1 was analysed by transient expression in 3 to 4 weeks old *N. benthamiana* leaves. The CDS was cloned in gateway pENTR/D-TOPO vector according to manufacturer’s protocol (K240020, Invitrogen, USA). The positive constructs were sequenced and further transferred to destination vector in-frame with superfold-green fluorescence protein (sGFP) in pGWB5 and pGWB6, tagged at carboxyl-and amino-terminal of HYL1 respectively, using Gateay LR Clonase II enzyme mix (11791-020, Invitrogen, USA). The recombinant pGWB5 and pGWB6 (encoding HYL1-GFP and GFP-HYL1) were finally transformed into *Agrobacterium* GV3101. The agroinfiltration was performed according to previously described (Raghuram *et.al.,* 2014). Briefly, the overnight grown culture was used for secondary inoculation in YEB broth and further incubated in shaker incubator at 28°C. The bacterial cells were pelleted down and washed with infiltration medium (10 mM MgCl2, 10 mM MES, pH 5.7, 150 l M acetosyringone) and final OD of the culture was maintained to 0.5 at A_600_. The culture was keep in dark for 2 to 3 hours. The culture was infiltrated on the lower epidermis of *N. benthamina* leaves by 1 ml needleless syringe. The fluorescence signal was monitored after 2 to 3 following day of post infiltration under a confocal laser scanning microscope to detect the sGFP fluorescence. To monitor the localization of GFP-HYL1 in the presence of AtMPK3, agrobacterium harbouring the AtMPK3 in pSPYCE(M) (AtMPK3-HA) (described by Raghuram et.al. 2014) were co-infiltrated in the leaves as above. The subcellular localized proteins were further prepared and analysed by subcellular fractions from the positive infiltrated leaves using Qproteome Cell Compartment kit (37502, QIAGEN). Immuno blotting were performed from subcellular enriched fractions by using anti-AtHYL1 and anti-GFP antibody (BB-AB0065, BioBharti, India) as described above.

### Multiple protein alignment analysis

To analyse the evolutionary relationship of dsRNA binding proteins (DRB1/HYL1) from *Arabidopsis thaliana* and other monocots and dicots by conserved dsRNA binding domain and putative MAP kinase sites. The protein sequences were downloaded from the National Centre for Biotechnology Information (NCBI) and Uniprot and multiple protein sequences were aligned by Uniprot align tool. First we aligned the HYL1 proteins from *Arabidopsis thaliana* with *Arabidopsis lyrata* (UniprotKB Identifier D7KJT2) and *Arabis alpine* (A0A087HMB7) and then with other close relative brassica members (*Brassica napus*/Q5IZK5, *Brassica oleracea*/A0A0D3DNR2, *Brassica_rapa*/M4EPS2). To further analyse the evolutionary basis of conservation of amino acid residues, we performed multiple sequence alignment of few monocots (*Populus trichocarpa*/ B9H6U2, *Vitis vinifera*/ A5BNI8-1, *Solanum lycopersicum*/ K4BU80) and dicots (*Zea mays*/B6TPY5-1, *Setaria italica*/ K3Y7A9, *Oryza sativa* subsp. *indica*/ I2DBG3, *Oryza sativa* subsp. *japonica*/ Q0IQN6, *Brachypodium distachyon*/ A0A0Q3F254, *Musa acuminate*/ M0RRC4).

### Prediction and evaluation of protein natural disordered regions

The AtHYL1 and RNA polymerase II largest subunit AtRPB1 Protein disorder were predicted using the VL-XT Predictor (Romero et al., 1997; 2001; Li et al., 1999) at PONDR (Predictor of Natural Disordered Regions; www.pondr.com) server. First, we predicted the natural disordered regions in the wild type proteins of both HYL1 and RPB1 protein followed by disordered enhanced phosphorylation throughout the protein then we substituted the putative MAP kinase target sites with phospho -null (T/S>A) or phospho mimetic (T/P>E/D) in the C-terminal of the both proteins followed by their disorder prediction. We further analysed the natural disordered regions in other members of AtDRBs (AtDRB2, AtDRB3, AtDRB4, AtDRB5) and OsDRBs (OsDRB1-1, OsDRB1-2, OsDRB1-3 and OsDRB1-4). The results were obtained in both tabular as well as the graphical data. The tabular data were further processed by using the GraghPad Prism software. The PONDR VL-XT predictor integrates three feedforward neural networks: the VL1 predictor (Romero et al. 1997), the N-terminus predictor (XN), and the C-terminus predictor (XC) (both from Li et al. 1999). VL1 was trained using 8 long disordered regions identified from missing electron density in x-ray crystallographic studies, and 7 long disordered regions characterized by NMR. The XN and XC predictors, together called XT, were also trained using x-ray crystallographic data, where the terminal disordered regions were 5 or more amino acids in length. Basically, the results are shown between 0 and 1 values. If the values for each residues exceeds or matches a threshold of 0.5 then the particular stretches are considered to be disordered.

### Prediction of Post-translational modification by Phosphorylation

The phosphorylation of AtHYL1 and AtRPB1 were predicted by disordered associated phosphorylation were predicted using Disorder Enhanced Phosphorylation predictor (DEPP), with protein sequence as input. The data were obtained in the graphical images and phosphorylation at serine, threonine and tyrosine were predicted when a target amino acid has >0.5 prediction value. The DEPP discriminates between the phosphorylation and non-phosphorylation sites by using the disorder regions around the target sites.

### Evolutionary relationship of AtHYL1 with DRB from other plants

The protein sequences were retrieved from the Uniprot database by BLAST search using AtHYL1 protein sequence as an input as described above. The phylogenetic tree was constructed using MEGA7 program for alignment of the sequences and construction of the phylogenetic tree by using maximum likelihood method based on the JTT matrix-based model. The tree with the highest log likelihood (-3606.13) is shown. Initial tree(s) for the heuristic search were obtained automatically by applying Neighbor-Join and BioNJ algorithms to a matrix of pairwise distances estimated using a JTT model, and then selecting the topology with superior log likelihood value. The tree is drawn to scale, with branch lengths measured in the number of substitutions per site. The analysis involved 14 amino acid sequences. All positions containing gaps and missing data were eliminated. There were a total of 244 positions in the final dataset. Evolutionary analyses were conducted in MEGA7.

## ACKNOWLEDGEMENTS

This work was supported by the core grant of National Institute of Plant Genome Research from the Department of Biotechnology (DBT), Government of India. PKB and RB are recipients of fellowship from the DBT, while DV from University Grant Commission, Government of India. We thank the Radioisotope facility, Confocal Microscopy Facility and the Central Instrumentation Facility of NIPGR, New Delhi, India.

## AUTHOR CONTRIBUTIONS

PKB and AKS designed the experiments and overall study. PKB performed all the experiments and wrote first draft of the manuscript. RB prepared the deletion constructs and *in-vitro* phosphorylation assay. DV cloned the localization and Y2H constructs and conducted part of Y2H assay. PKB, RB and AKS participated in the discussion and progress of work. PKB, RB wrote the manuscript. Final draft was supervised by AKS.

---

**Supplemental figure 1.**
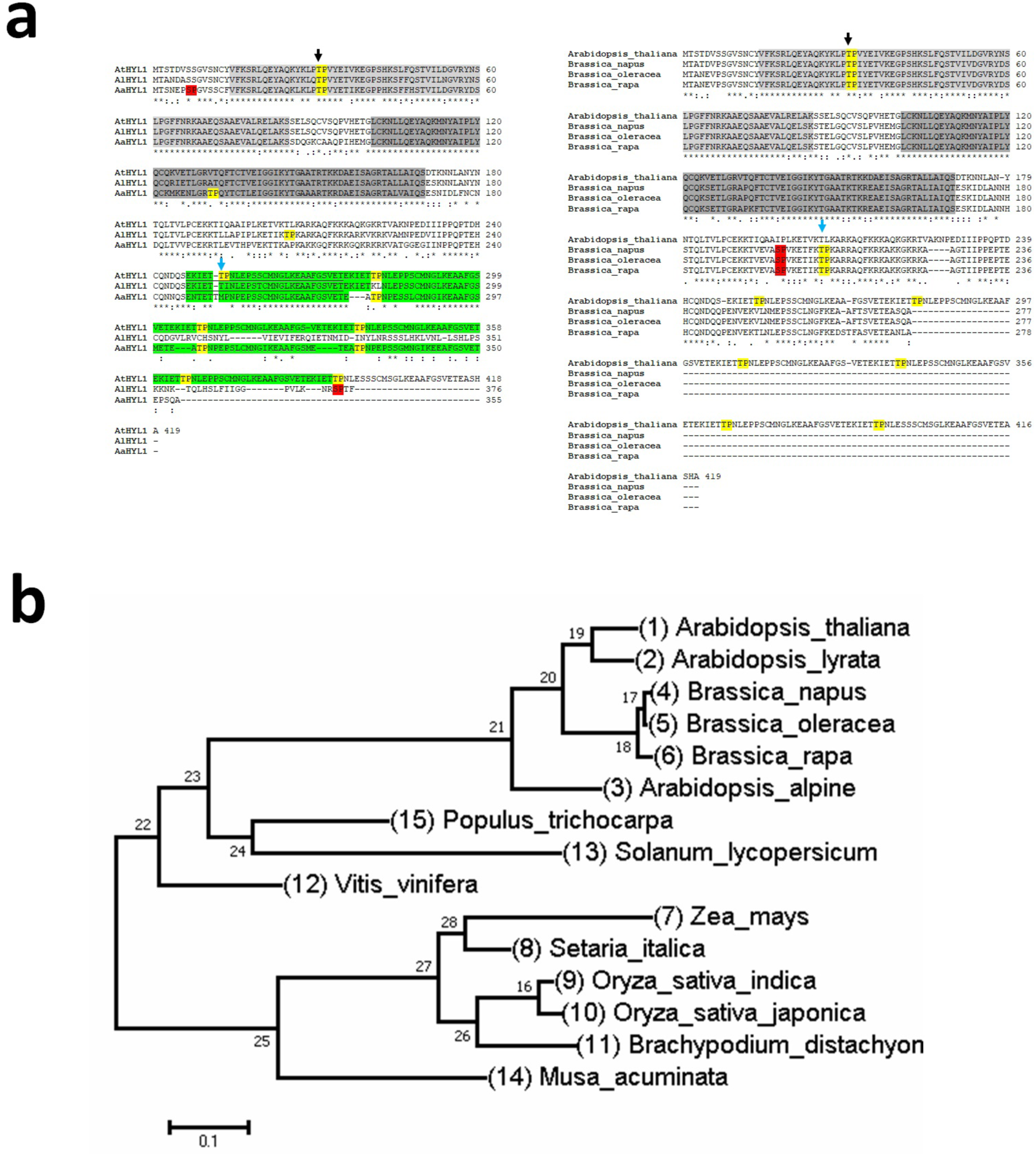
Multiple protein sequence alignment and evolutionary analysis of AtHYL1 and DRBs from plant kingdom. a, (Left side) Protein alignment of HYL1 from *A. thaliana* (At), *A. lyrata* (Al) and *A. alpine* (Aa). The C-terminal repeats (28a.a.) are highlighted by green. The (right side) images represents the protein alignment of Arabidopsis close relative brassica members. The conserved dsRBD-I and dsRBD-II are highlighted by light and dark grey respectively. The conserved MAP kinase sites are highlighted and pointed by black head arrow and newly emerged TP motif by sky blue arrow head. **b**, the phylogenetic tree was constructed by using Maximum Likelihood method based on the JTT matrix-based model using MEGA7 and representative image is shown.

**Supplemental figure 2.**
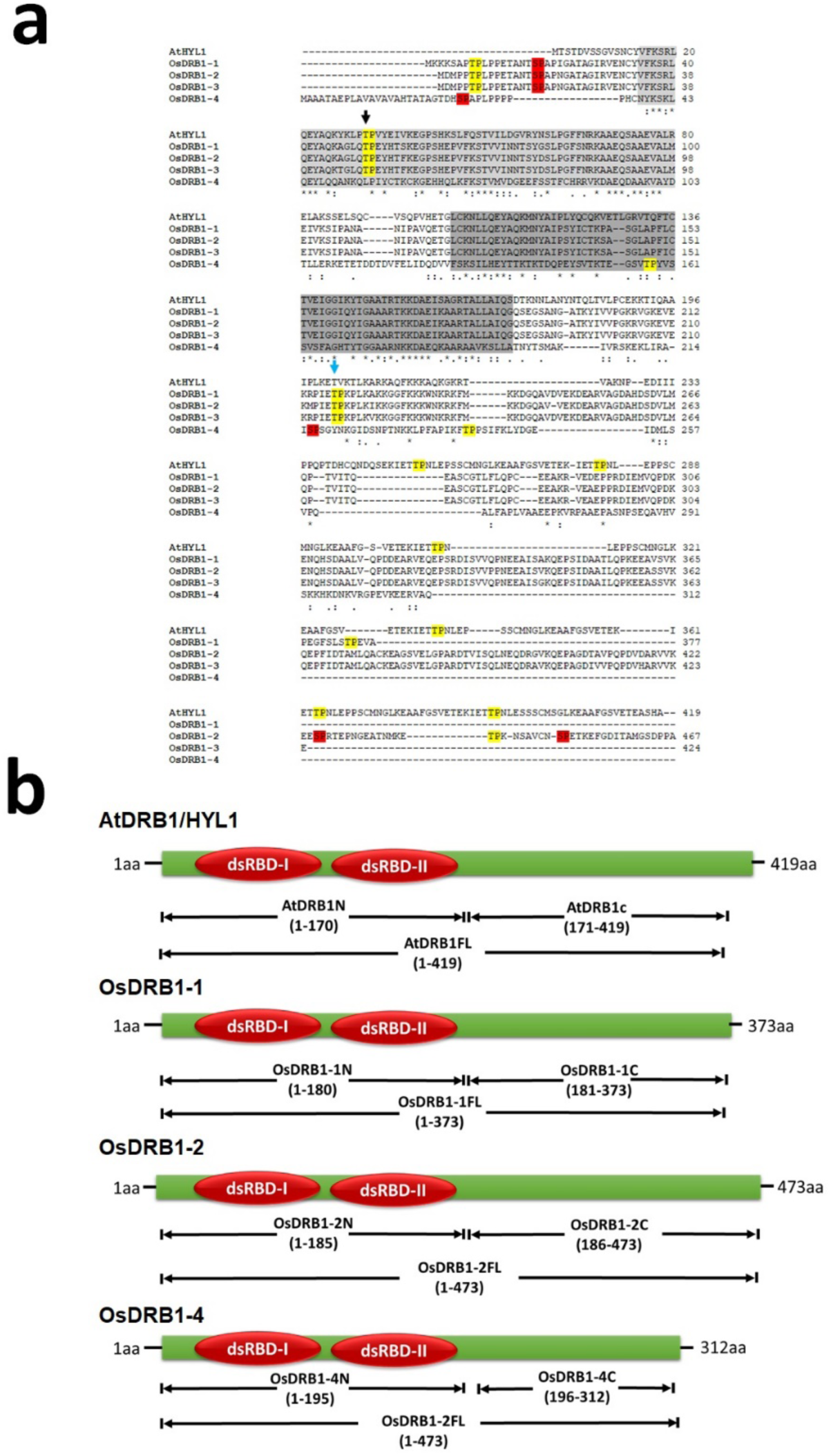
**a.** Protein alignment of AtHYL1, OsDRB1-1, OsDRB1-2, OsDRB1-3 and OsDRB1-4. **b**. schematic representation of deletion constructs of OsDRBs and AtHYL1 for protein expression and in-vitro phosphorylation assay.

**Supplemental figure 3.**
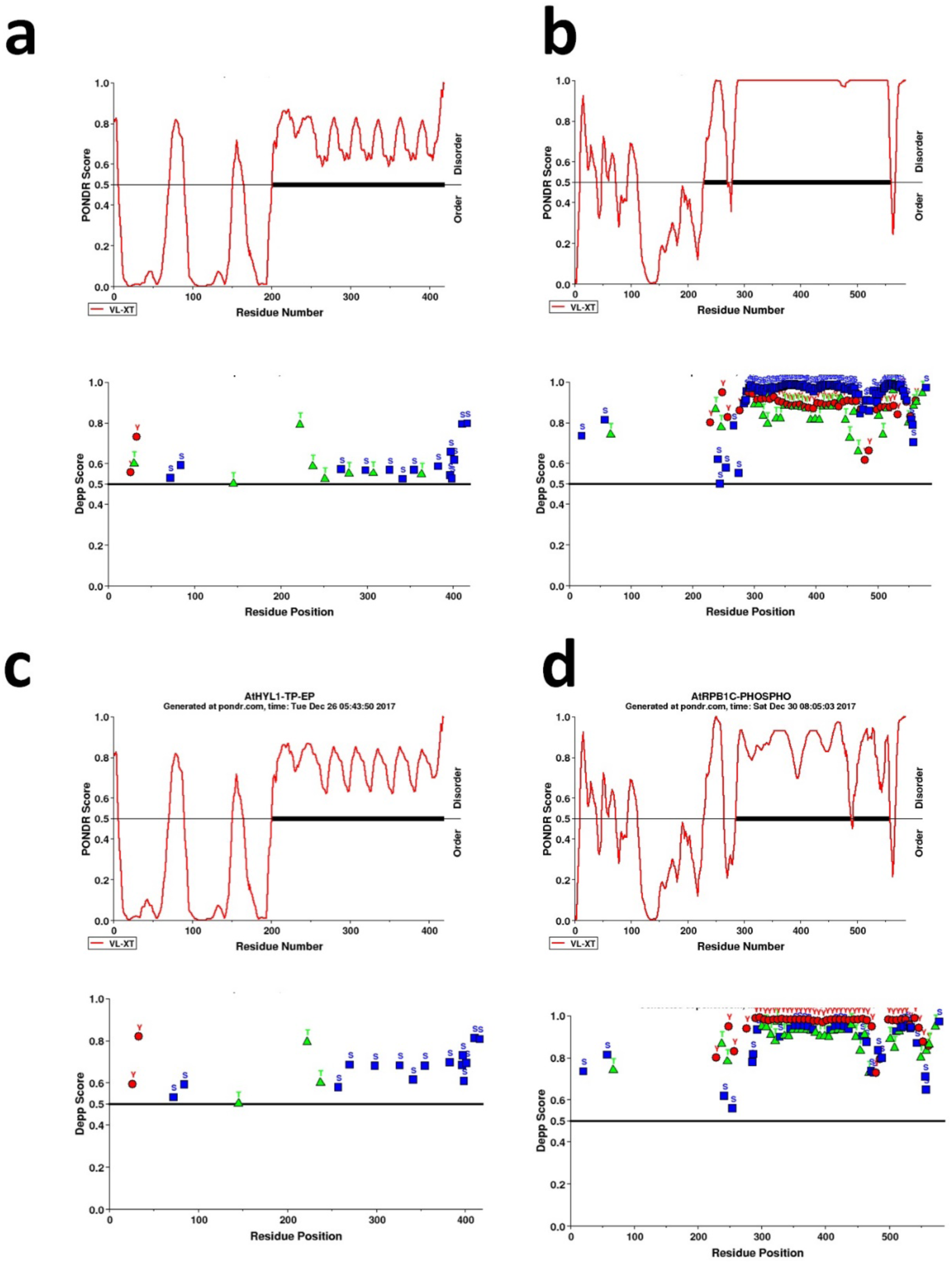
**a** & **b.** Prediction of natural disordered regions in AtHYL1(AT1G09700) full length and AtRPB1(AT4G35800) C-terminal (1254 to 1839 amino acids) upper images and disorder enhanced phosphorylation lower images. **c** & **d.** Substitution of serine and threonine to aspartic acid or glutamic acid in the putative MAP kinase sites to mimic as phosphorylated isoform and their subsequent analysis as above. The procedure is described in the method sections.

**Supplemental figure 4.**
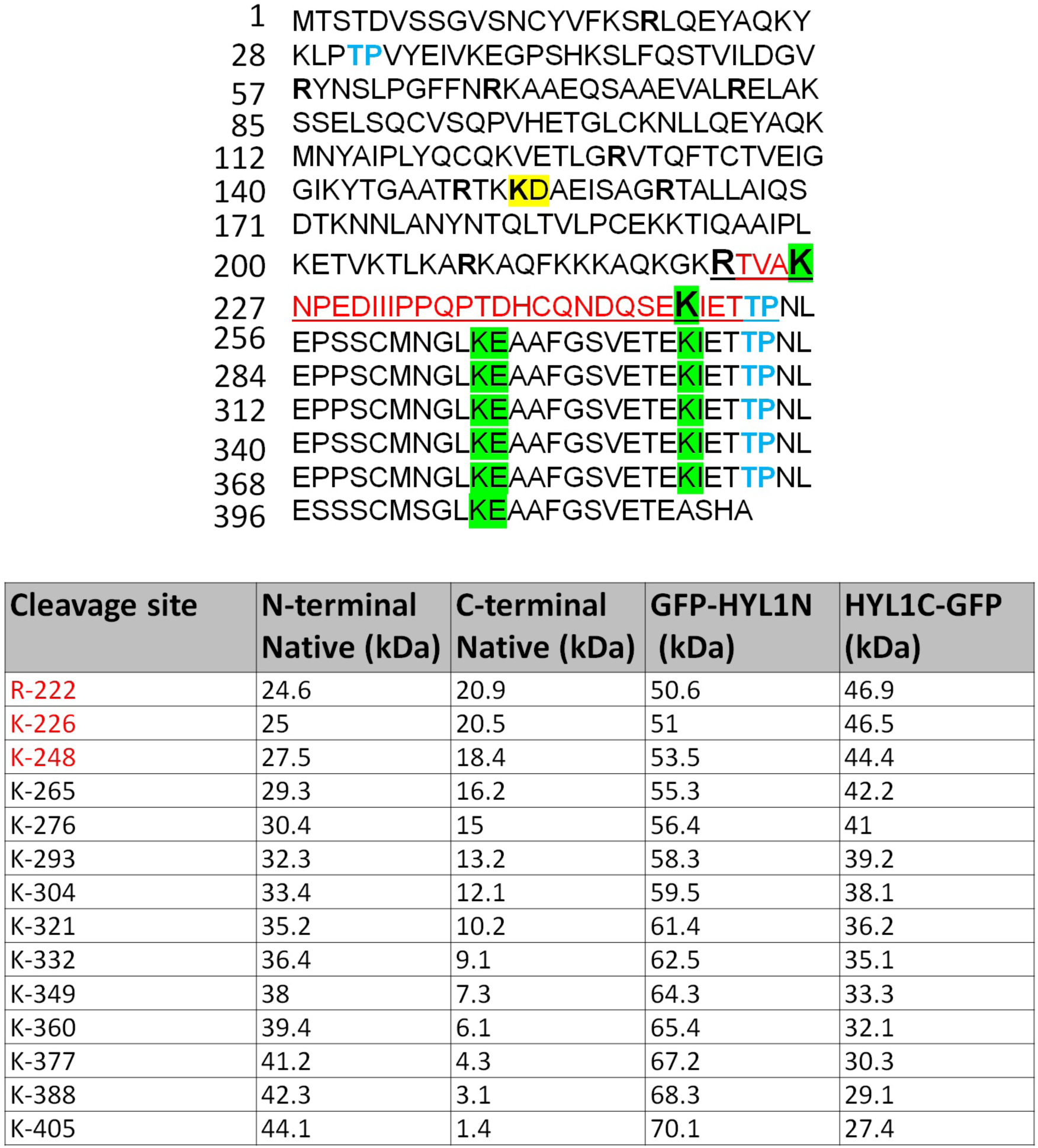
Properties of HYL1 protein sequence. Representation of amino acid sequence and post-translational modification and putative trypsin cleavage sites. The putative bipartite NLS is underlined and coloured red. The putative MAP kinase sites coloured by sky blue and C-terminal multiple cleavage sites are highlighted by green. The most probable site for trypsin cleavage in the NLS are bold and larger in size. The size of different cleaved products of HYL1 after proteolysis is predicted based on the site of cleavage and represented their expected molecular weight with or without GFP for in-vivo localisation and western blotting purpose.

**Supplementary table 1:**
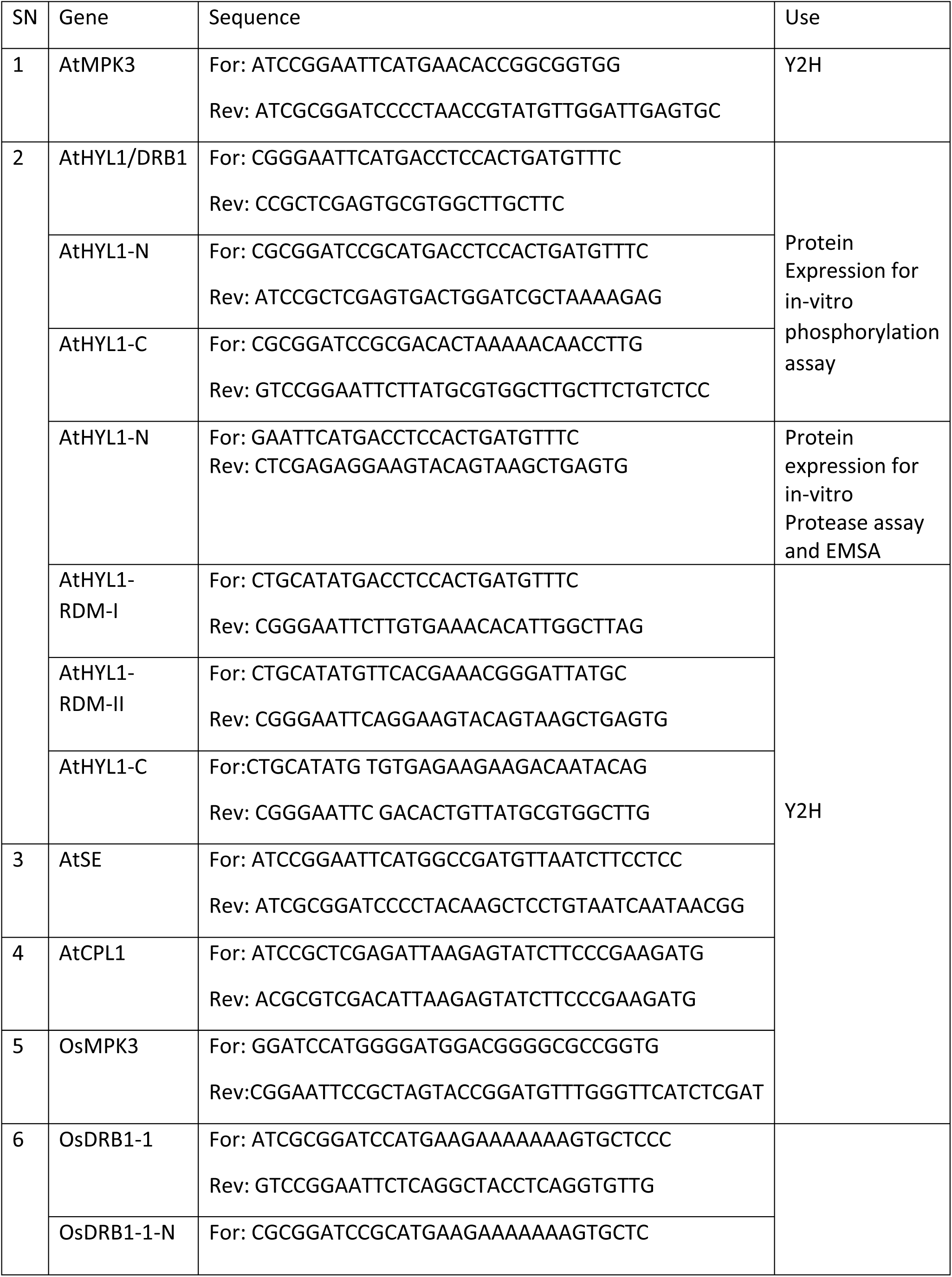
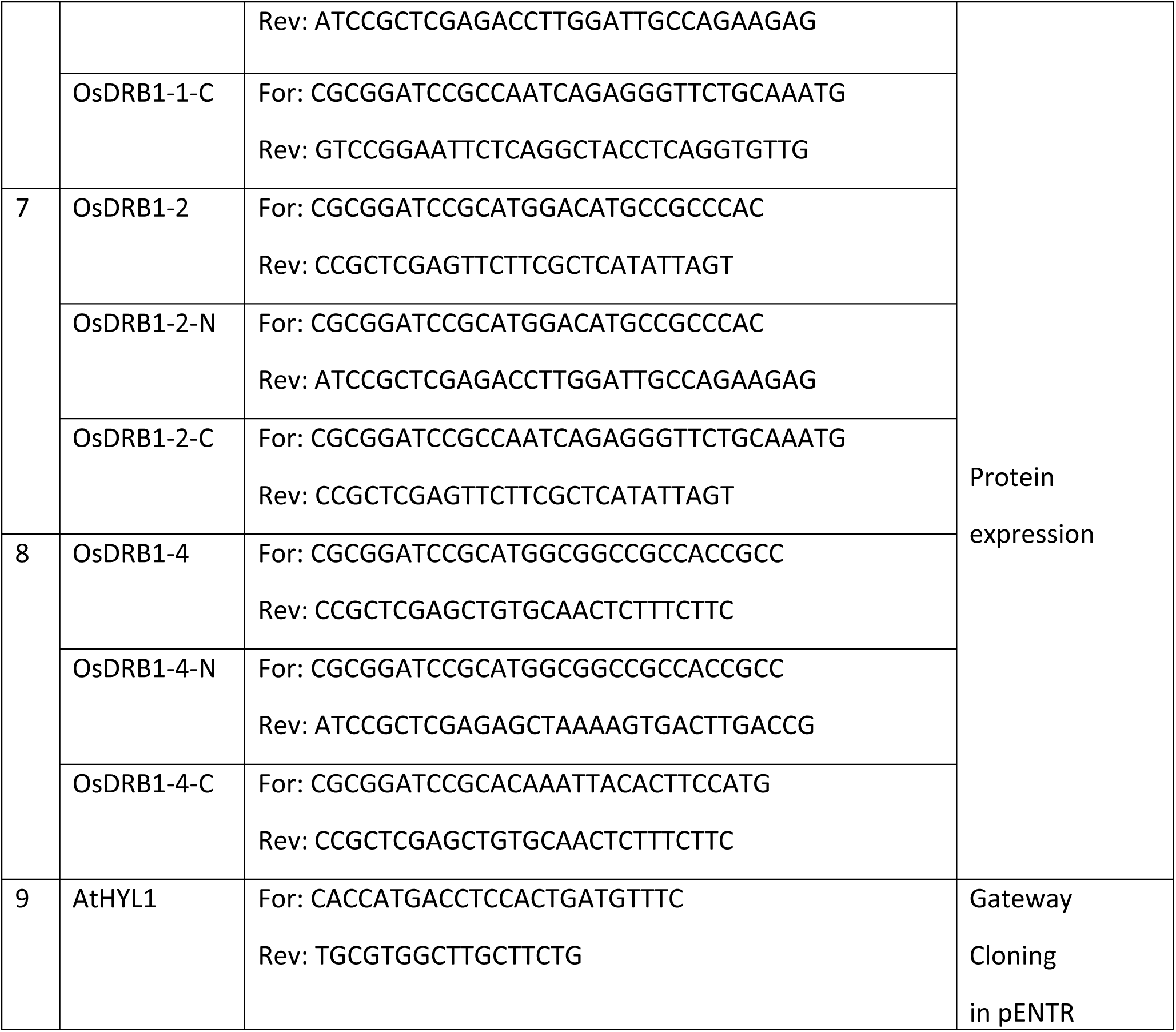
List of primers used in the present study

